# The Use of Informativity in the Development of Robust Metaviromics-based Examinations

**DOI:** 10.1101/054635

**Authors:** Siobhan C. Watkins, Thomas Hatzopoulos, Catherine Putonti

## Abstract

The field of metagenomics has developed insight into many of the complex microbial communities responsible for maintaining life on this planet. Sequencing efforts often uncover novel genetic content; this is most evident for viral metagenomics, in which upwards of 90% of all sequences demonstrate no sequence similarity with present databases. For the small fraction which can be identified, the top BLAST hit is often posited as being representative of the phage taxon. However, as previous research has shown, the top BLAST hit is sometimes misinterpreted. Furthermore, the appearance of a particular gene homolog is frequently not representative of the presence of the particular taxon in question. To circumvent these limitations, we have developed a new method for the analysis of metaviromic datasets. BLAST hits are weighted, integrating the sequence identity and length of alignments as well as a phylogenetic signal. A genic rather than genomic approach is presented in which each gene is evaluated with respect to its information content. Through this quantifiable metric, predictions of viral community structure can be made with greater confidence. As a proof-of-concept, the approach presented here was implemented and applied to seven metaviromes. While providing a more robust means of evaluating metaviromic data, the tool is versatile and can easily be customized to investigations of any environment or biome.

## Background

Bacterial viruses (bacteriophages) play an essential role in shaping microbial populations. They drive community structure through the mediation of mortality, and shape diversity - fundamentally - through their role as agents of genetic mobility (Wilhelm & Suttle, 1999; Canchaya et al., 2003; Beredjeb et al., 2011; Clokie et al. 2011; Winget et al., 2011; Brum et al. 2016). Their impact has been described at higher trophic levels (Rohwer & Thurber, 2009; Jover et al., 2014); phages affect microbial processes on a global scale. In addition to their influence in the environment, evidence has uncovered that phages can contribute to human disease (e.g. Holmes 2000) and may play a role in human health as part of the human microbiome (e.g. Willner et al., 2012). Whole genome sequencing (WGS) inquiries of complex viral communities (metaviromics) have been pivotal in ascertaining both the ubiquity of phages as well as the sheer number of phages on Earth (Edwards & Rohwer, 2005). As such, a wide variety of environments have been probed, from the world's oceans (Hurwitz & Sullivan, 2013) to extreme environments (Gudbergsdottir et al., 2015); from deserts (Fancello et al., 2013), to the human gut (Minot et al., 2013).

In contrast with cellular organisms, no conserved coding regions are ubiquitous among all viral species. Efforts to utilize genes coding for structural proteins have given limited insight into the diversity of defined communities of phages (Dorigo, Jacquet & Humbert, 2004; Wilhelm et al., 2006). Similarly, DNA polymerases have been used as markers for specific groups of phages (Breitbart, Miyake & Rohwer, 2004). However, the study of viral communities based on the examination of whole genomes is widely considered to be the most robust approach to exploring phage diversity in the environment. The approach taken for analyzing WGS data sets within metaviromics has paralleled that of metagenomics of bacterial and archaeal populations - reads or contigs are compared to known, characterized sequences within public data repositories. Although a powerful tool, the generation of metaviromic surveys, a literal “who’s who” of the communities present, is confounded by bioinformatic challenges unique to the examination of phages. Currently, only a small fraction of the genetic diversity that phages represent is characterized - and it is certainly likely that the large gaps in our knowledge define key processes. However these general gaps are translated directly from the genome level; most characterized phages contain a surfeit of genes for which there are no known homologs (Hatfull, 2008). In addition, the current collection of characterized genomes is sparse; presently, there are just over 2000 phage genomes deposited in RefSeq, and strains that infect laboratory bacterial models are overrepresented. Therefore, phages represent a remarkable reservoir of undiscovered genetic diversity (Suttle, 2007).

For the few viral species which can be identified, typically via BLAST searches, the single best hit is often posited as being representative of the phage taxon containing the homologous region: a method employed by many metagenome studies and analysis tools (e.g., Huson & Weber, 2013; Wommack et al., 2012; Keegan et al., 2016; Roux et al., 2014). This approach, however, can be misleading; genes present within annotated phage genomes may not be true indicators of the phage species. For instance, such genes may be bacterial in origin (e.g. Mann et al., 2003; Thompson et al., 2011; Thompson et al., 2011; Lindell et al., 2005; Gao, Gui & Zhang, 2012). Thus hits to such genes would be indicative of either bacterial DNA within the sample sequenced or acquisition of the bacterial genome (which need not be exclusive to the taxa represented in the sequence data repositories). In a recent metaviromic survey of the nearshore waters of Lake Michigan, further investigation of viral species with the most “hits” revealed that the matches were localized to a particular gene(s) within the genome, and therefore indicative of the presence of a specific gene rather than that of the species (Watkins et al., 2015). Moreover, as was the case with one of these phages - Planktothrix phage PaV-LD, BLAST results were indicative of the presence of bacterial genes. Several Planktothrix phage genes exhibit greater sequence similarity to bacterial proteins rather than other phage sequences (Gao, Gui & Zhang, 2012). Over half of the publicly available datasets in the viral metagenomic sequence web server MetaVir (Roux et al., 2014) include hits to this phage (including samples unlikely to harbor the phage's cyanobacterial host species), indicating that misreporting is widespread. Thus, a “BLAST and go” approach for species identification must be replaced by a more rigorous assessment of each individual BLAST hit result.

Herein we present a new, quantifiable, method for assessment of BLAST results, in an attempt to address the aforementioned challenges. This approach can be applied to all studies, regardless of the niche under investigation, as sequence similarity to databases is weighted. Weighting takes into consideration not only the sequence identity between the metavirome contig and the database record, but also the length of the alignment, and more importantly the informativity of the match. This latter metric captures the taxonomic signal within sequence similarity results. Thus, a species' presence or absence within a population can be determined with greater confidence. As a proof-of-concept, we examined seven publicly available freshwater DNA metagenomic datasets.

## Materials and Methods

*Viral gene datasets*. Sequence data were retrieved from NCBI in January 2016. For the analysis of Pbunalikeviruses, amino acid and nucleotide sequences for the Pbunalikeviruses Pseudomonas phage PB1 (Accession Number: NC_011810) and Burkholderia phage BcepF1 (Accession Number: NC_009015). All phage nucleotide sequences (omitting those belonging to the Pbunalikevirus genomes listed in Supplemental Table 2) were retrieved through an advanced search via the NCBI website with the following query: PHG[Division] NOT (txid538398[Organism] AND …) in which the list of Pbunalikeviruses were removed from the search by their taxonIDs (as indicated by “…”). In total over 500000 individual records were retrieved.

*Metaviromic datasets*. SRA records were collected from the SRA database. Supplemental Table 1 lists all of the datasets included in the proof-of-concept study. Each SRA record (line listed in the Supplemental Table 1) was considered as an individual sample. (Note, two samples are aggregates of more than one SRA record, both belonging to Metavirome IV, as they were combined in the downloadable file from SRA.) Each individual sample was next assembled using Velvet (Zerbino & Birney, 2008) with a hash size of 31. PB1 protein sequences were directly compared to these assembled contigs, rather than raw reads, via blastx.

**Table 1.** Freshwater DNA metaviromic studies retrieved from NCBI’s SRA database.

| Metavirome | Environmental Niche | Number of Samples | Sequencing Technology | Mbp Total | Reference |
| --- | --- | --- | --- | --- | --- |
| I | Lake Michigan nearshore | 40 | Illumina | 6 909 | Watkins et al., 2015; Sible et al., 2015 |
| II | Lake Bourget | 2 | 454 | 698 | Roux et al., 2012 |
| III | Kent SeaTech tilapia pond | 3 | 454 | 47 | Dinsdale et al., 2008 |
| IV | Lake Limnopolar | 2 | 454 | 18 | López-Bueno et al., 2009 |
| V | Reclaimed water samples | 6 | 454 | 364 | Rosario et al., 2009 |
| VI | Lake Ontario | 3 | 454 | 223 | n/a |
| VII | Feitsui Reservoir | 5 | 454 | 86 | Tseng et al., 2013 |

## Results and Discussion

### Determination of Informativity Metric for Quantifying Hits

***Establishing a Phylogenetic Signal Threshold*.** To ascertain the presence/absence of specific taxon within a metagenome, we suggest a threshold to differentiate between informative and uninformative hits. The phylogenetic signal threshold *T* is determined through a two-step process prior to evaluation of the metagenomic data. Firstly, for a given taxon of interest, each annotated coding region is compared to all annotated sequences within the genome of a known relative. Thus, each coding region's sequence *x* (*x*ϵ*X*, where *X* is the set of sequences for all coding regions annotated within the genome of the taxon of interest) is compared to each coding region's sequence *g* (*g*ϵ*G*, where *G* is the set of sequences for all coding regions annotated within the genome of a known relative). The use of a known relative genome establishes if and how conserved the coding region is between known, related strains/species. Where sequence homology is detected, the sequence identity and query coverage of the match is recorded: *S_1_* and *Q_1_*, respectively.

In the second step, each coding region’s sequence is compared again, this time to the sequences for all annotated coding regions for the group assayed by the metagenomic study (e.g. phages, all viruses, bacteria, archaea, etc.), however, those belonging to the phylogenetic group containing the taxon of interest and the known relative considered in step one are omitted. Many hits may be recorded for a particular gene *x*. Thus the best hit, both with respect to the sequence identity and the query coverage of the match, is selected; *S_2_* and *Q_2_* denote this best match's sequence identity and query coverage, respectively. A phylogenetic signal threshold *T* is defined as *T*={*S_1_*−*S_2_*, *Q_1_*−*Q_2_*} where the subscripts 1 and 2 represent the sequence identity and query coverage of the match detected from steps one and two, respectively. Figure 1 illustrates the two-step process, the *T* values produced.

It is important to note, that the phylogenetic group used for comparison is user defined. For instance, in order to ascertain if a gene can be used to distinguish between the presence/absence of a particular species, one may consider the phylogenetic group to be inclusive only of strains of the species. Therefore in this case, the most distant relative belonging to the phylogenetic group in step one would be the closest related species. If a more distant relative, say the most distantly related species of the same genus, were to be investigated, then the phylogenetic signal threshold *T* would serve as a means to distinguish between the presence/absence of a subset of the species (inclusive of the taxon of interest) within the genus. This flexibility enables the researcher to define and control the granularity of his/her analyses. In addition to the intended purpose of establishing the phylogenetic signal threshold, the two-step process can provide insight into putative horizontally acquired elements and gene loss events within a phylogenetic group. For example, instances in which the gene did not include a homolog in the most distant relative but did exhibit sequence similarity to a gene within the genome of another phylogenetic group. Furthermore, the two-step process can identify genomes which have been taxonomically misclassified - such instances would result in high *S_2_* and *Q_2_* scores for a large majority of the genes.

**Figure 1.**
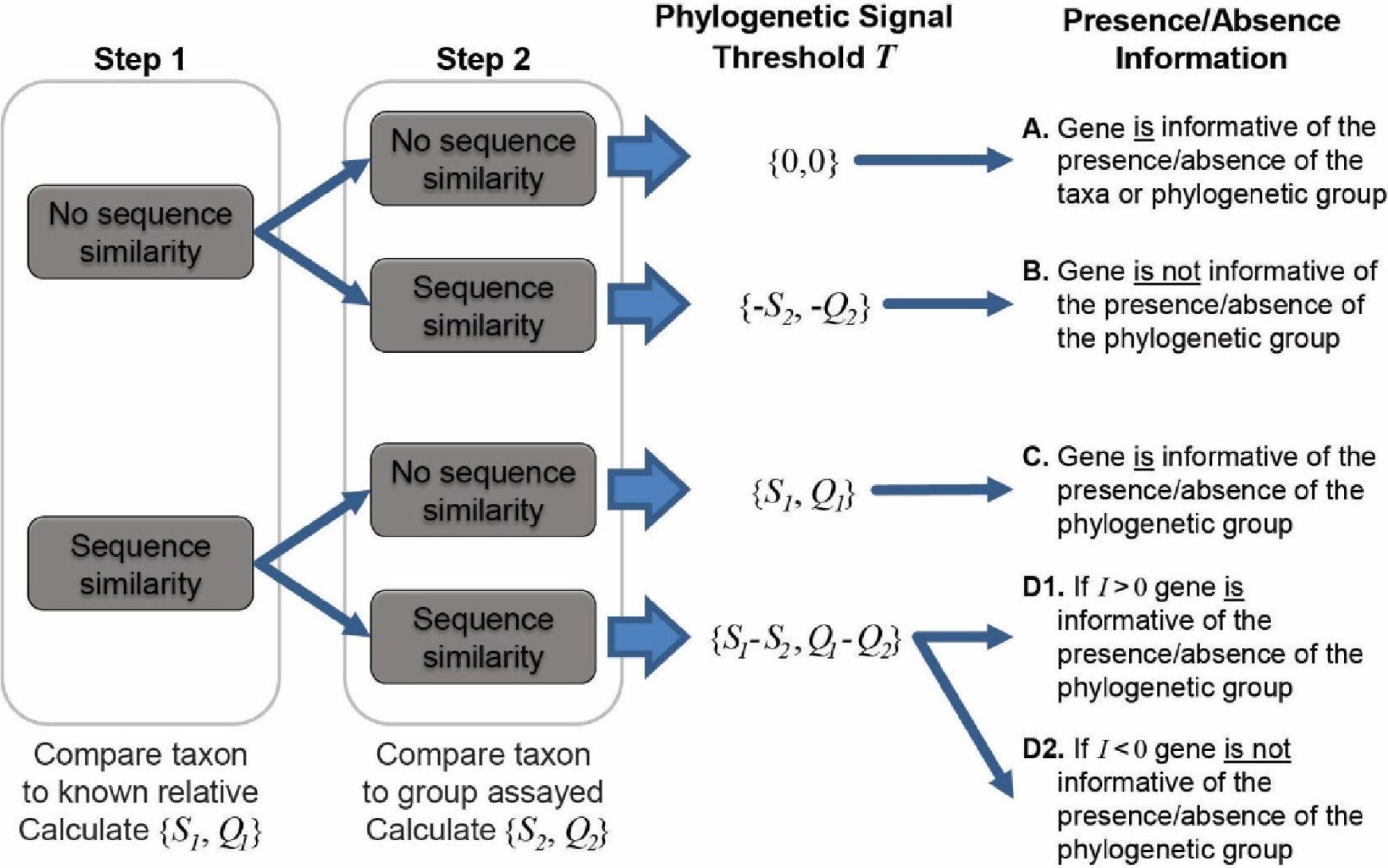
Two-step process for determining the phylogenetic signal threshold *T* and the information which can be gained regarding the presence/absence of a taxon’s phylogenetic group. *S_1_* and *S_2_* represent the sequence identity of homologies identified in step 1 and 2, respectively. Likewise, *Q_1_* and *Q_2_* refer to the query coverage of the match detected in step 1 and 2, respectively.

***Using Informativity to Ascertain Confidence in OTU Calls*.** As indicated in Figure 1, when the set *T* is greater than or equal to zero (outcomes A, C, and D1), the presence of a specific gene can provide insight. OTU calls are informed by this threshold to decipher BLAST analyses of metaviromic datasets as some hits may be to genes which are conserved and thus poor indicators of a species’ or taxa’s presence or absence. For a given “hit” within a metaviromic dataset, the sequence identity and query coverage, *S_H_* and *Q_H_* respectively, is assessed relative to the phylogenetic signal threshold *T* for the gene producing the match. Genes in which *T* < 0 have already been classified as uninformative (Figure 2). Now hits which fall below the gene’s threshold, {*S_H_*, *Q_H_*}−*T* < 0, are also classified as uninformative. Hits which are above the threshold are considered informative. The informativity *I* of each hit is quantified based upon deviation from this threshold *T* such that *I*= {*S_H_*, *Q_H_*}−*T*. *I* can range from 0 (equivalent to the threshold *T*) to 100 (*T*={0,0}, *S_H_* = 100% sequence identity and *Q_H_* = query coverage of the gene). Thus genes with a large value of *I* are strong indicators of the presence of a particular taxon.

**Figure 2.**
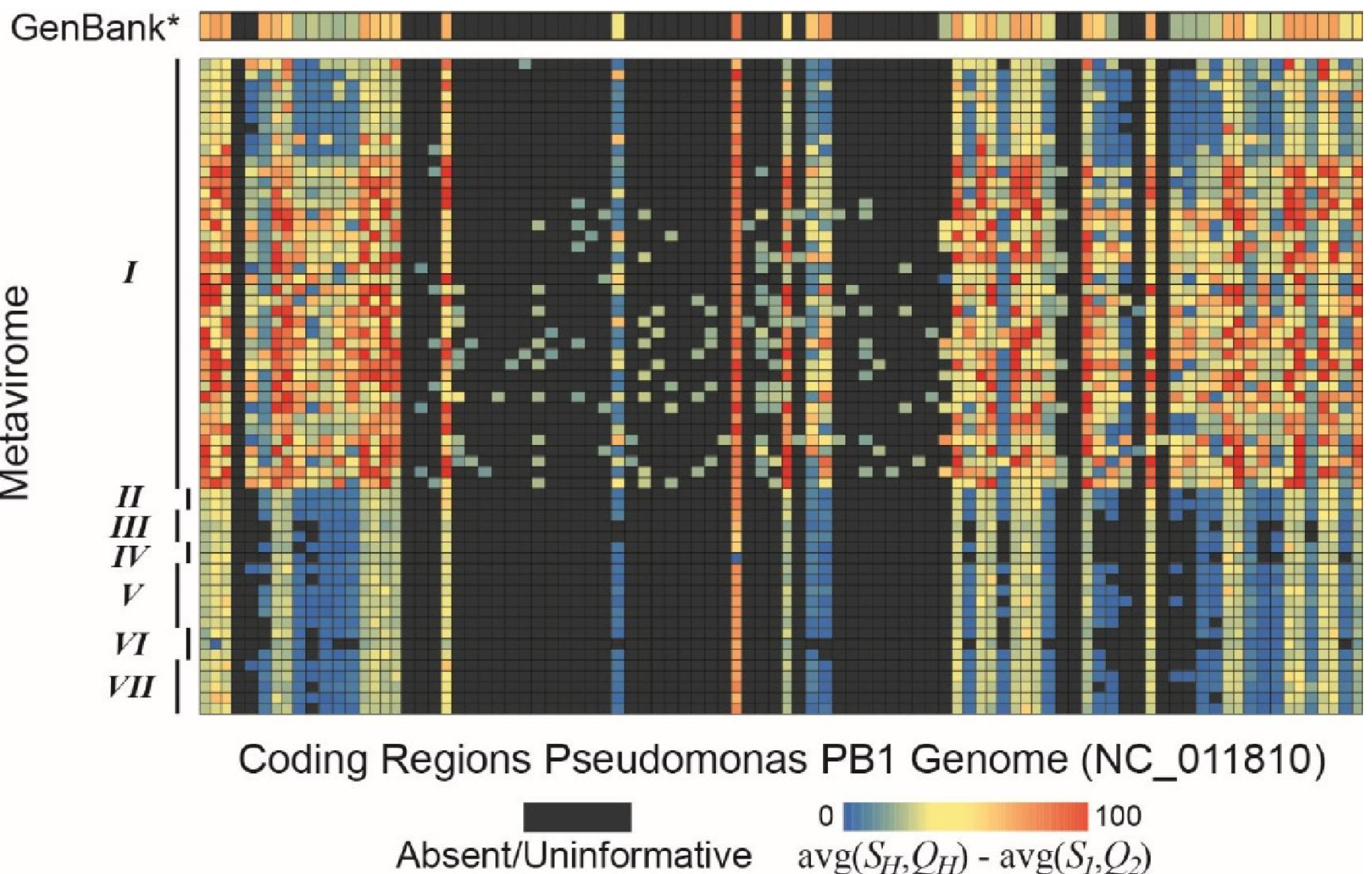
BLAST hits to PB1 genes within both the set of non-Pbunalikevirus viral genomes and seven freshwater DNA metaviromic datasets (Table 1). Hits (*S_H_* and *Q_H_*) are qualified relative to the sequence similarity shared between PB1 and its distant Pbunalikevirus relative, Burkholderia phage BcepF1 (*S_1_* and *Q_2_*).

### Implementation

The posited method for assessing the informativity of metagenomic hits was implemented using a series of BLAST databases and BLAST searches. First, a collection of all coding regions (either nucleotide or amino acid sequences) were retrieved for the taxon of interest (*X*) as well as all genes annotated within the user defined genome of the selected relative (*G*). A local BLAST database was created for *G*, and the genes belonging to *X* were queried against the local database. The sequence identity and query coverage of the match detected for the best hit for each gene was then parsed from the BLAST results quantifying each gene’s *S_1_* and *Q_1_* values. Next, a BLAST database was created using all characterized, annotated sequences other than those associated with the phylogenetic group. Each of the genes for the taxon of interest *X* was queried against the second local database; the results were again parsed for each gene’s *S_2_* and *Q_2_* values so that the phylogenetic signal threshold *T* could be calculated.

A metagenomic dataset was next evaluated, comparing each read or contig against a collection of annotated gene sequences. While we implemented this step locally, users with limited computational resources can utilize a resource such as MG-RAST (Keegan, Glass & Meyer, 2016), MEGAN (Huson & Weber, 2013), VIROME (Wommack et al., 2012), or MetaVir (Roux et al., 2014) and use the remotely generated BLAST results produced for further analysis here. Each BLAST hit was next assessed with respect to its scores {*S_H_*, *Q_H_*} relative to that of the gene’s threshold *T*. Informative results were written out to file, including the values of *I*, *T*, and{*S_H_*, *Q_H_*}. The user can then evaluate the likelihood of a particular taxon or phylogenetic group’s presence within the metagenomic sample based upon the *I* values for informative genes, as described. Taking into consideration the number of informative genes detected within a metgenomic sample and their individual *I* values can leverage additional confidence in calling OTUs.

The described process has been automated via a Python script and calls to commands within the BLAST+ command line application. Users must supply or specify the fasta format files for the taxon of interest (*X*), the genome of a known relative (*G*), and the group assayed (less the taxonomic group of interest). If metagenomic comparisons are to be conducted locally, the user must also supply the metagenomic dataset. The script has been designed for both ease of use as well as flexibility, such that analyses can be tailored to the environmental niche and/or hypothesis under investigation. Most importantly, this script is a light-weight solution which can be integrated into the standard method of metaviromic analyses. The script and documentation are publicly available through http://www.putonti-lab.com/software.html.

### Proof-of-Concept

Our group previously isolated and characterized phages similar to the Pseudomonas phage PB1 (Malki et al., 2015), therefore we sought to examine populations of PB1 within other freshwater environments. Thus, each gene annotated for the PB1 genome (Accession Number: NC_011810) (Ceyssens et al., 2009) was compared first to the set of genes for the most distant relative of PB1 within its genus Pbunalikeviruses, Burkholderia phage BcepF1 (Accession Number: NC_009015). For each gene the *S_1_* and *Q_1_* values were computed. Next each gene annotated for the PB1 genome was compared via blastx to all genes from viral species other than those annotated as Pbunalikevirus in GenBank (see Methods), determining the values of *S_2_* and *Q_2_*.

Surprisingly the majority of the PB1 genes exhibited greater sequence similarity to sequences within this collection than they did to the Burkholderia phage BcepF1. This led us to manually inspect the genomes producing these hits. In doing so, we identified a number of viral strains assigned to the taxonomic level of “unclassified Myoviridae” within NCBI, rather than “Pbunalikeviruses”. These genomes were thus removed from the collection of non-Pbunalikevirus viral gene sequences (as they are in fact Pbunalikeviruses) and blastx was run again. (See Supplemental Table 2 for a list of the genomes reclassified here as Pbunalikeviruses.) Threshold *T* was then calculated for all 93 annotated PB1 genes. This threshold is visually represented in Figure 2 in the row marked as “GenBank*”. This variation is represented as a single measure, the average of *S_H_* and *Q_H_* (*S_2_* and *Q_2_* in this case) less the average of *S_1_* and *Q_1_*. Here we can see that several gene sequences (as indicated by the color scale) had better “hits” to records within the GenBank collection queried than they did to the Burkholderia phage BcepF1; gray blocks signify that no or weaker homology was detected (*T*≤0).

The methodology developed here was then applied to seven freshwater DNA metaviromic studies (Table1); a list of the SRA datasets from each study is provided in Supplemental Table 1. Reads from all seven metavirome datasets were first assembled (see Methods for details). The contigs were then compared to the PB1 genome via blastx. Figure 2 graphically represents these results. Again, each gene’s best hit within each metavirome sample was qualified (colored) with respect to its value relative to *S_1_* and *Q_1_*. From Figure 2, one can readily identify that not all genes provide an equal signal as to the presence or absence of PB1 within the sample, some serve as better markers. For instance, there are several genes which have a greater sequence similarity to the PB1 genome than PB1 has to BcepF1; these hits are represented within the heatmap. However non-Pbunalikevirus phage sequences may exhibit equivalent or greater sequence similarity to the PB1 gene sequence (as shown in the GenBank* row). The informativity metric provides a quantifiable confidence in assigning the presence/absence of a taxon. Thus, the informativity *I* of each BLAST hit within the metaviromic samples was calculated. In doing so, individual genes which provide a strong phylogenetic signal for the Pbunalikeviruses can readily be identified. Figure 3 represents the results of this computation, in which each hit to a PB1 gene is now assessed in light of the phylogenetic signal.

In an effort to assess the strength of the metric presented here, we evaluated the raw BLAST results of the datasets and a BLAST score-based analysis. The BLAST results of Metaviromes II, IV, V, and VII are publicly available through the web service MetaVir (Roux et al, 2014). Nine of the samples from Metavirome I are also available through MetaVir. It is important to note that in contrast to the uniform method in which the metavirome samples were preprocessed here (see Methods), the sequences submitted to MetaVir may be assembled or raw sequences. Furthermore, MetaVir conducts BLAST comparisons against the RefSeq viral database, whereas here we have included all partial and complete phage sequences from GenBank which is several magnitudes of difference greater in size. Nevertheless, hits to the Pbunalikeviruses (Supplemental Table 2) genomes were identified in all five MetaVir datasets; the Lake Michigan and Lake Bourget samples (nine samples from Metavirome I and both samples from Metavirome II) produced the most BLAST hits to the Pbunalikeviruses genomes (hundreds to thousands). As MetaVir determines taxonomy based upon the best BLAST hit, these best hits were next evaluated. All five datasets again included hits which were classified as Pbunalikeviruses.

As Figure 3 shows, Metavirome I (the Lake Michigan metaviromes generated by our group (Watkins et al., 2015; Sible et al., 2015)) identifies many informative genes indicative of the presence of Pseudomonas phage PB1. Metaviromes II, V, and VII contain informative hits to 1, 2, and 1 PB1 genes respectively. Their informativity, however, is low, i.e. {*S_H_* and *Q_H_*} ≈ *T*. This would suggest that PB1 is not present within the sample: rather a homolog of the gene is present. The prevalence of informative genes within several of the samples of Metavirome I and the lack thereof in the other metaviromes suggests that PB1 and likewise other Pbunalikeviruses are not present (or at the least not prevalent) in the other metaviormes. As viral sequence databases expand through the isolation and characterization of additional viral strains, the threshold *T* is likely to change thus providing greater confidence in the evaluation of BLAST hits for OTU calling.

**Figure 3.**
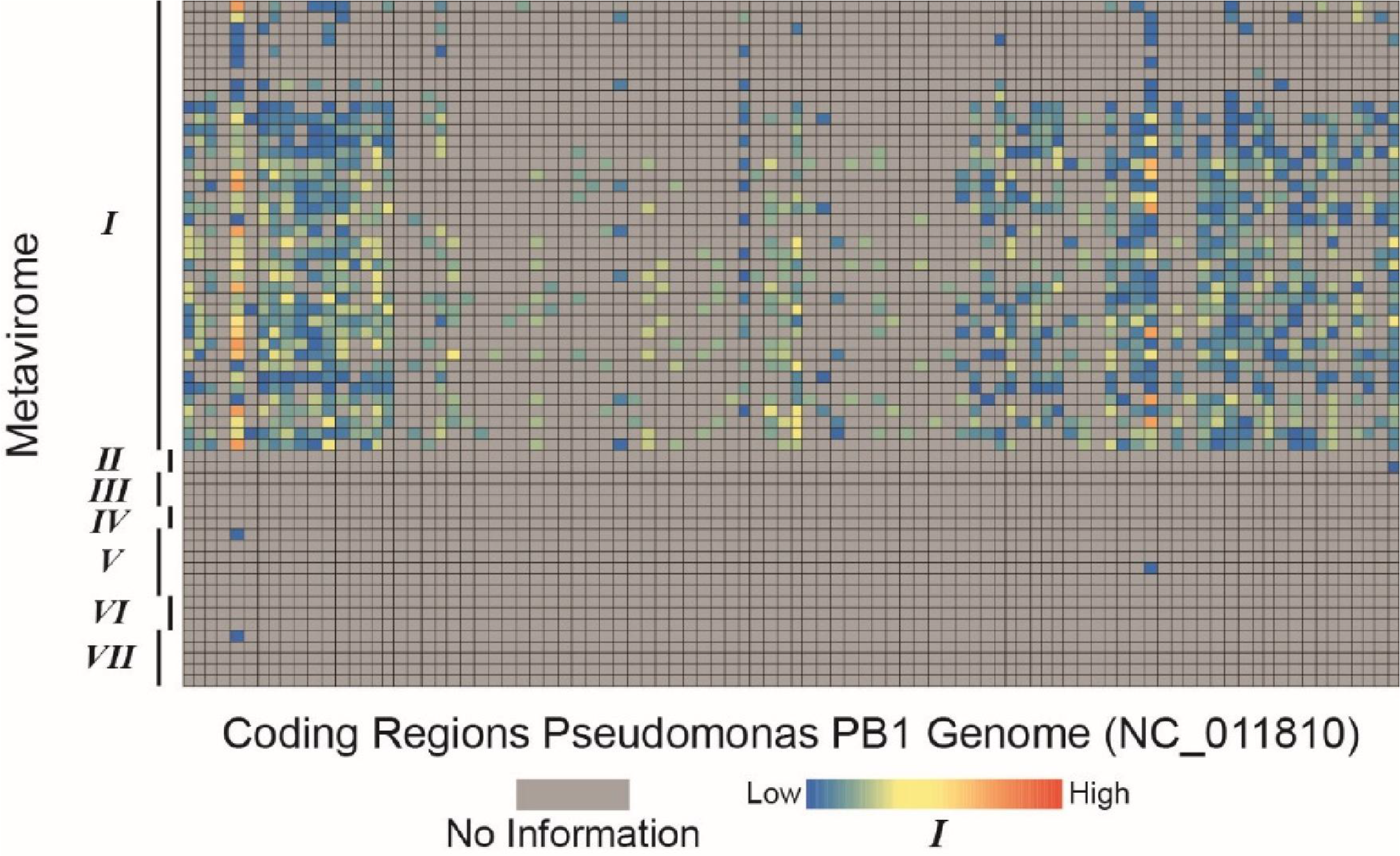
Informativity of hits to PB1 genes within seven freshwater DNA metaviromic datasets (Table 1).

### Conclusions

The presented method for extrapolating the presence/absence of microbial taxa is both robust and versatile. Although specifically developed to tackle some of the challenges facing metavirome studies, it can be applied to any WGS dataset. Specifically, the proof-of-concept investigation of seven freshwater metavirome datasets can be applied in the effort to identify novel strains and species of phages with
confidence. Many of the prokaryote members of the human microbiome are undergoing examination, but exploration of human viromes is the next frontier (Abeles & Pride, 2014; Ogilvie & Jones, 2015). As such, these studies will face many of the same challenges that are detailed as part of the presented study. Nevertheless, improved bioinformatic tools for mining metaviromic analyses, coupled with further physical isolation and characterization of viral species have the potential to greatly expand our knowledge of the viral diversity on Earth.

## Acknowledgements

The authors would like to thank Ms. Katherine Bruder, Alexandria Cooper, Kema Malki, and Emily Sible for their contributions to general research investigating Pbunalikeviruses.

## Supplemental Data

**Supplemental Table 1.**
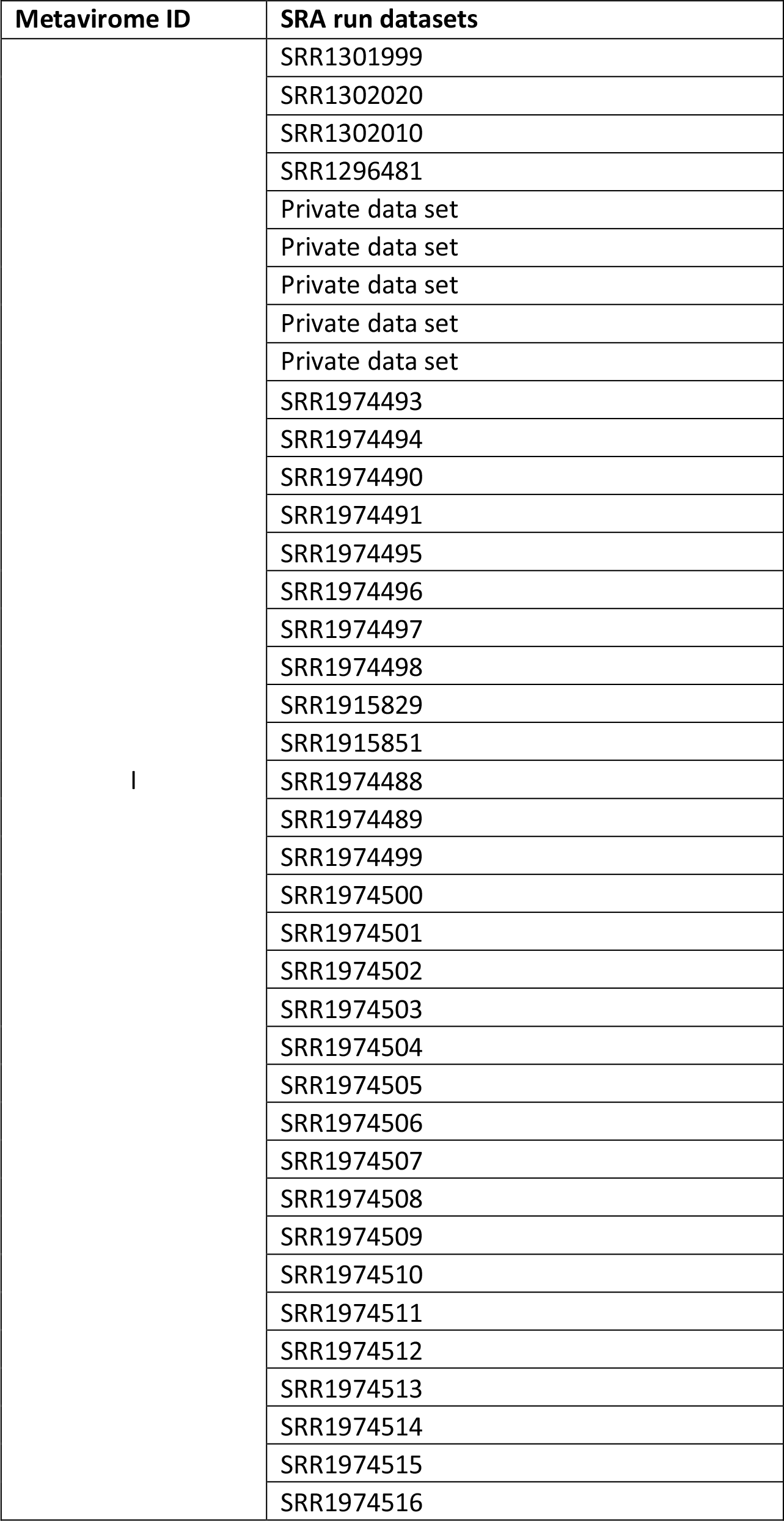
SRA datasets from each study.

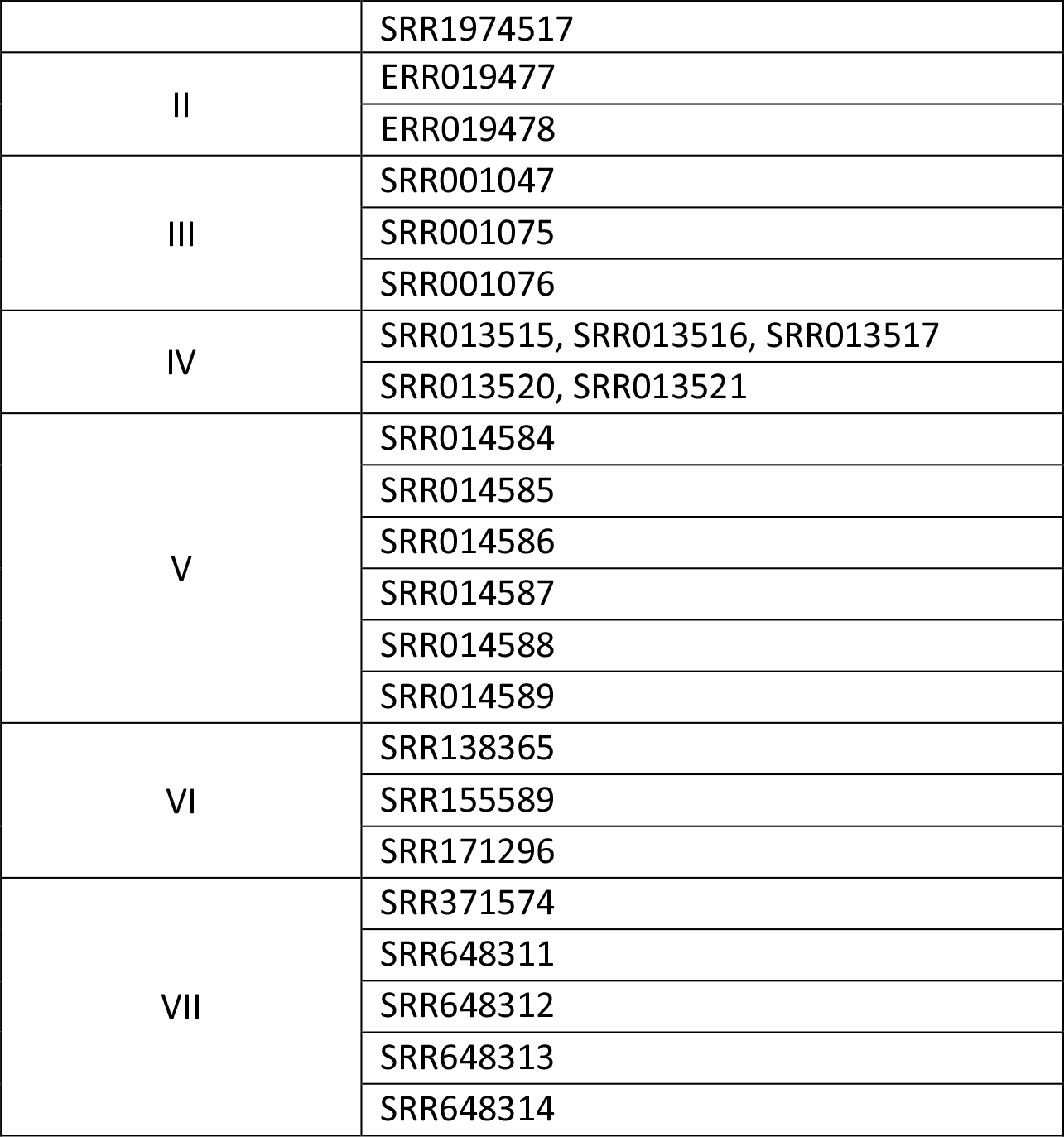

**Supplemental Table 2.** Pbunalikevirus genomes

| Phage | Genome size | NCBI Assigned Taxonomy | GenBank Accession No. |
| --- | --- | --- | --- |
| PB1 | 65764 | Pbunalikevirus | NC_011810 |
| SN | 66390 | Pbunalikevirus | NC_011756 |
| 14-1 | 66238 | Pbunalikevirus | NC_011703 |
| LMA2 | 66530 | Pbunalikevirus | NC_011166 |
| LBL3 | 64427 | Pbunalikevirus | NC_011165 |
| F8 | 66015 | Pbunalikevirus | NC_007810 |
| BcepF1 | 72415 | Pbunalikevirus | NC_009015 |
| PaMx13 | 66450 | Pbunalikevirus | JQ067083 |
| pp DL52 | 65867 | Pbunalikevirus | KR054028 |
| pp SPM-1 | 65729 | Pbunalikevirus | NC_023596 |
| pp vB_PaeM_C1-14_Ab28 | 66181 | unclassified Myoviridae | NC_026600 |
| pp DL60 | 66103 | unclassified Myoviridae | KR054030 |
| pp KPP12 | 64144 | unclassified Myoviridae | NC_019935 |
| pp NH-4 | 66116 | unclassified Myoviridae | NC_019451 |
| pp vB_PaeM_PAO1_Ab27 | 66299 | unclassified Myoviridae | NC_026586 |
| pp vB_PaeM_PAO1_Ab29 | 66326 | unclassified Myoviridae | LN610588 |
| pp JG024 | 66275 | unclassified Myoviridae | NC_017674 |
| pp DL68 | 66111 | unclassified Myoviridae | KR054033 |
| S12-1 | 66257 | unclassified Myoviridae | LC102730 |
| R18 | 63560 | unclassified Myoviridae | LC102729 |

